# Regulatory plasticity of the human genome

**DOI:** 10.1101/2024.11.13.623439

**Authors:** Jaya Srivastava, Ivan Ovcharenko

## Abstract

Evolutionary turnover in non-coding regions has driven phenotypic divergence during past speciation events and continues to facilitate environmental adaptation through variants. We used a deep learning model to identify the substrates of regulatory turnover using genome wide mutations mimicking three evolutionary pathways: recent history (human-chimp substitutions), modern population (human population variation), and mutational susceptibility (random mutations). We observed enhancer turnover in approximately 6% of the whole genome, with more than 80% of the novel activity arising from repurposing of enhancers between cell-types. Frequency of turnover in a cell-type is remarkably similar across the three pathways, despite only ∼19% overlap in the source regions. The enhancers predisposed to turnover display reduced evolutionary constraints and are depleted in GWAS variants. Most of these occur within 100kb of a gene, with the highest turnover occurring near neurodevelopmental genes including CNTNAP2, NPAS3, and AUTS2. Flanking enhancers of these genes undergo high turnover irrespective of the mutational model pathway, suggesting a high plasticity in neurocognitive evolution. Based on susceptibility to random mutations, these enhancers were identified as vulnerable by nature and feature a higher abundance of cell type-specific transcription factor binding sites (TFBSs). Our findings suggest that enhancer repurposing within vulnerable loci drives regulatory innovation while keeping the core regulatory networks intact.

## 1. Introduction

Rapid evolution of enhancers between species has put them in the spotlight of evolutionary mechanisms that have driven the course of metazoan evolution (García-Pérez et al., 2021; Villar et al., 2015; Weiss et al., 2021). For instance, comparative epigenomics approaches have identified changes within enhancers regulating anatomical, neurodevelopmental and immunological features that contribute to evolution of human-specific phenotypes (Andrews et al., 2023; Cotney et al., 2013; Prescott et al., 2015; Uebbing et al., 2021; Villar et al., 2015; Xue et al., 2023). Species-specific regulatory landscapes evolve largely due to changes within unconstrained enhancers that are susceptible to environmental interactions and lead to the origins of species biased features (Andrews et al., 2023; Cotney et al., 2013). Gene expression is more tolerant to variations in regulatory activity due to robustness conferred by enhancer redundancy which mitigates the impact of changes in a single enhancer (Kvon et al., 2021; Wong et al., 2015). However, even closely related species like humans and chimpanzees with ∼98% genomic identity show significant differences in morphology, immunity, and cognition, largely due to enhancer divergence, highlighting their role in reshaping regulatory landscapes (Prescott et al., 2015; Vermunt et al., 2016).

Reduced evolutionary constraints in the non-coding genome, or lack thereof in some regions of the genome, has led to the accumulation of millions of variants in the human population (1000 Genomes Project Consortium, 2010). These variations enable the evolution of novel enhancers either *de novo* by creating Transcription Factor Binding Sites (TFBSs) in neutral DNA sequences or through exaptation of existing enhancers for novel regulatory activity (Long et al., 2016), which we refer to herein as enhancer repurposing. This leads to spontaneous gains and losses in enhancer activity, i.e., enhancer turnover which creates inter-individual phenotypic variations (Castelijns et al., 2020). Over time, adaptive changes in enhancer function culminate in evolution of lineage-specific regulatory novelty (Long et al., 2016). Potential impact of these variants over time have led to emergence of traits like lactose tolerance (Bersaglieri et al., 2004), skin pigmentation (Lamason et al., 2005), malaria resistance (Kwiatkowski, 2005), etc., all of which have evolved as adaptations to environmental factors within very short timescales.

However, rapid turnover within enhancers begets questions concerning the regulatory plasticity of our genome. For instance, what fraction of the genome undergoes regulatory turnover because of accumulating variants? What is the most common mechanism by which enhancers acquire novel activity in short timescales? Are certain cell-types, genes or TFBSs more amenable to changes than others? How do the paths of regulatory reprogramming that have occurred in the past compare with the potential changes that will come into effect with variants that are accumulating in our genome? Methodological challenges hinder large-scale analysis of evolutionary dynamics driven by variants. In this study, we address these questions using a deep learning (DL) model that can quantify the effect of genome wide variants on enhancer activity.

A significant proportion of genomic variations within humans and between humans and closely related primates is attributed to single nucleotide variants (SNVs) (Byrska-Bishop et al., 2022; Waterson et al., 2005). Despite over 98% genome similarity between humans and their closest relatives, chimps, enhancer divergence explains key phenotypic differences between the two species (Waterson et al., 2005). The substantial presence of orthologous genomic regions has simplified comparative epigenomics in identifying phenotypic differences due to enhancer divergence (Prescott et al., 2015; Whalen & Pollard, 2022). We used these genome-wide differences as a metric to define an evolutionary step leading to species-biased phenotypes. Subsequently, we generated model genomes by introducing chimp-specific variants, population variants, and random variants into the human reference genome to quantify and compare regulatory changes that occur under distinct mutational landscapes. Our results show that 80% of novel enhancer activity arises from repurposing existing enhancers, altering activity between cell-types. Enhancer repurposing leads to species- and cell-type specific gene expression profiles and provides insights into the role of non-cognate cell-types in complex diseases. We also extended our approach to generate ten random genomes and identified evolutionarily constrained robust enhancers which are resistant to mutations and encode transcription factor binding site (TBFS) profiles that are distinct from mutation prone enhancers. Our results support a model of regulatory innovation within cell-type specific enhancers that accommodate adaptive changes while preserving the core regulatory network that is driven by cell ubiquitous evolutionarily constrained enhancers.

## 2. Results

### Quantifying the impact of sequence variation on enhancer activity using deep learning

As a first step towards predicting the impact of genomic variants on enhancer activity, we used a previously developed DL model dubbed TREDNet that could predict the effects of non-coding variants on enhancer activity (Hudaiberdiev et al., 2023; S. Li et al., 2023). This two-phase DL model with convolutional neural network (CNN) framework was demonstrated as a successful approach in prioritizing causal variants of type 2 diabetes (Hudaiberdiev et al., 2023) and in identifying mutations associated with human cognition (S. Li et al., 2023). We used chromatin accessibility profiles from ATAC-Seq or DNAse-Seq overlapping with H3K27ac histone marks as a proxy for active enhancers to train cell-type specific models for 67 ENCODE defined biosamples (Methods). Area under the receiver operating characteristic curve (auROC) ranged from 0.9 to 0.98, and the area under the precision-recall curve (auPRC) ranged from 0.56 to 0.85 for the cell-types. Difference in TREDNet score of a sequence fragment before and after introduction of variant(s) was used as a predictor of differential enhancer activity. Sequences with scores below and above the threshold (set at FPR_1%_ score) before and after introducing variants, respectively, were termed as gain of activity (GoA) enhancers. Loss of activity (LoA) enhancers were identified using the same approach with opposite criterion.

To validate the use of TREDNet scores in predicting GoA and LoA enhancers, we used experimental results for differential H3K27ac peak intensities in humans and chimps, which have been shown to correlate with differential gene expression between species and lead to species biased phenotypes (Prescott et al., 2015; Whalen et al., 2023). A recent study profiled H3K27ac marks in chimp and human Neural Progenitor Cells (NPCs) in an attempt to identify human accelerated regions involved in neurodevelopment (Whalen et al., 2023). We used peak intensities of H3K27ac marks from this study as a proxy for enhancer activity in human and chimp NPCs. Specifically, we identified 8,304 orthologous regions with H3K27ac marks in both species and leveraged differential peak intensities arising from sequence variants as indicators of species-specific enhancer activity. We scored these regions with a TREDNet model trained for NPCs using sequences from human (hg38) and chimp reference genome (PanTro6) and compared H3K27ac signal intensities with our model’s predictions. Our model predicted 135 regions as human GoA enhancers. These regions are validated by significantly higher human H3K27ac intensities (Wilcoxon p-value < 0.001, Figure S1) as compared to 1,217 regions for which are predicted as conserved by the DL model and do not reveal any differences in H3K27ac signals. These human-chimp specific NPC enhancer GoA observations in addition to the previously reported several other experimental lines of evidence (D. Huang & Ovcharenko, 2024; Hudaiberdiev et al., 2023; S. Li et al., 2023) validate our model’s capability to identify variants that disrupt, strengthen, and/or introduce enhancer activity.

### Fate of cis-regulatory landscape differs across different cell types in the face of genome wide variants

We next leveraged our model’s predictions to quantify the genome-wide impact of regulatory variants on enhancer activity within the modern human population and potential future genome variants. On average, the genome of every individual varies by ∼0.4% of which a majority comprise SNVs (“A Map of Human Genome Variation from Population Scale Sequencing,” 2010). We were thus interested to know if these population variants, as well as an evolutionarily unconstrained set of random variants, have the potential to drive regulatory changes as significant as those linked to SNVs between closest primate relatives, humans and chimps. To this extent, we first generated a model chimp genome by introducing 36,621,296 single nucleotide substitutions specific to chimp (PanTro6) into the human (build 38) reference genome while omitting other types of sequence variation (see Methods for details). Next, we generated a model 1000g genome by introducing combined variants from the 1000 Human Genomes project (1000g) catalog (Fairley et al., 2020). Finally, we constructed a series of synthetic genomes containing the same number of variants as the number of SNVs separating humans and chimps introduced at random locations in the human genome. All model genomes contain the same number of variants as the number of human-chimp substitutions to replicate a timescale that culminates in a possibly distinct phenotype. The model chimp genome, 1000g variants genome and random variants genome are henceforth referred as model chimp, 1000g and random genome, respectively. Between the three model genomes, 5% of the variants shared between the model chimp and 1000g genome, and 1% between the other two pairs (Figure S2). However, they do not impose significant biases in turnover between the model genomes (Supplementary note 1).

The 1000g genome contains a significant proportion of rare variants with minor allele frequency (MAF) <= 0.01. Rare variants have been proposed to have a larger impact on gene expression (Cavalli et al., 2016; Momozawa & Mizukami, 2021). However, we observed that allele frequency does not impact the key conclusions drawn in this study from enhancer turnover observed across cell-types and model genomes (Supplementary note 2).

Next, we quantified enhancer turnover in 67 cell-types across the three model genomes. TREDNet scores for 1kb tiled genome windows (3,088,221 regions in total) in the three model genomes were compared with the human reference genome using cell-type specific TREDNet models and GoA and LoA enhancers were identified using whole genome as a template, excluding promoter-proximal and coding regions (Figure 1A). The combined set of GoA and LoA enhancers from all cell-types in a model genome collectively spans approximately 6% of the whole genome. The proportions of both GoA and LoA enhancers vary significantly across cell-types, with the most enhancer gains and losses in the stomach and least in the thyroid gland (Figure 1B). Thus, cell-types like thyroid glands, and spleen are the most robust unlike stomach, myotube, and fibroblasts which undergo high rates of turnover. Additionally, the numbers of GoA and LoA enhancers in each cell-type is invariable across the three model genomes, even though they do not frequent the same loci, as we observed only ∼19% overlap between the GoA or LoA enhancers of the three model genomes. In addition, the number of GoA enhancers are equivalent to that of LoA enhancers regardless of the model genome and the cell-type (Figure 1B). Together, these observations indicate the presence of redundant loci that compensate for the gain and loss of enhancer function and preserve the total complement of functional enhancers in different mutational landscapes.

**Figure 1.**
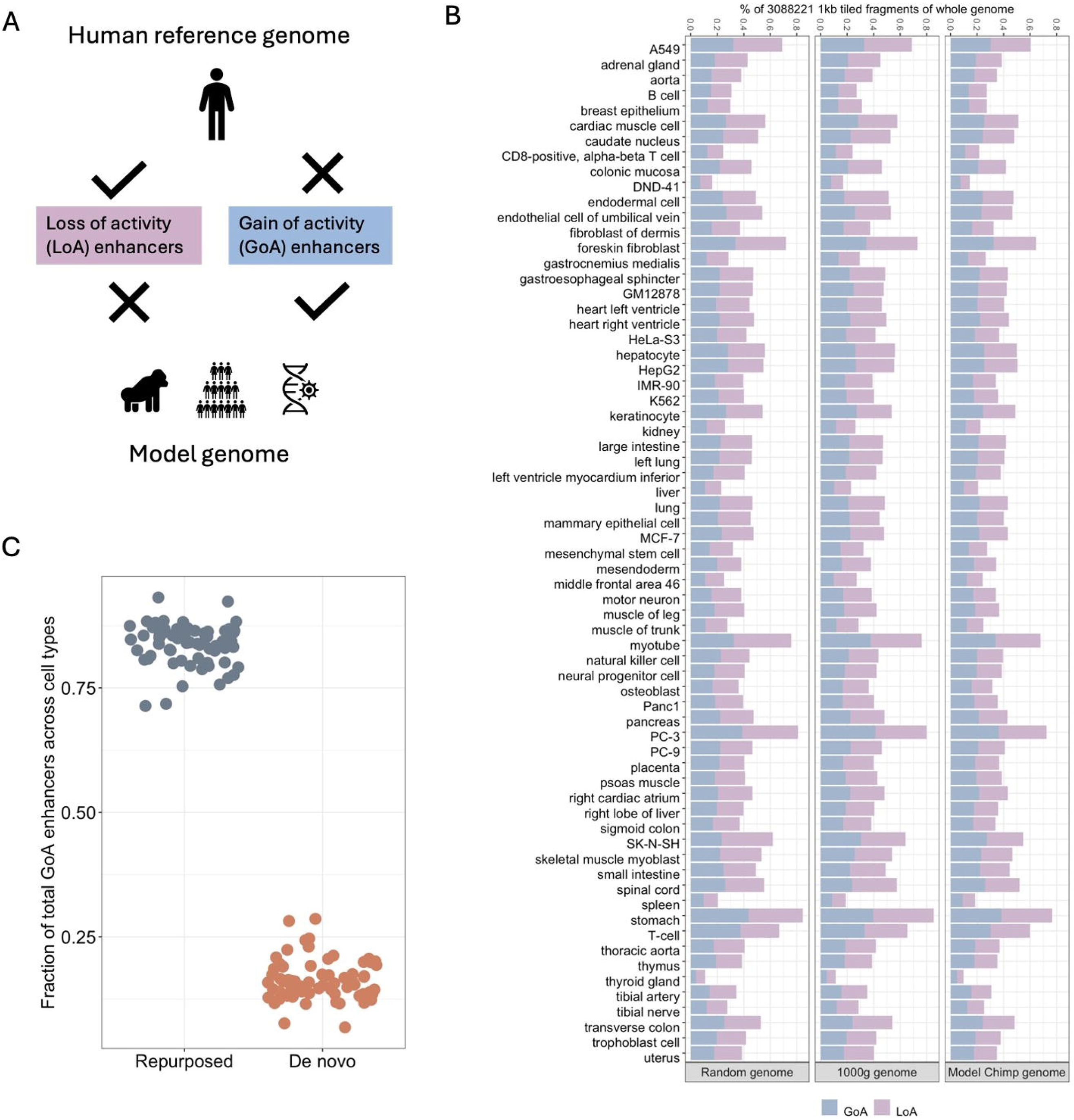
GoA and LoA enhancers in the three model genomes. (A) Identifying GoA and LoA enhancers. If a genomic region is inactive in human reference genome but active in a model genome, it is a GoA enhancer of the model genome. Conversely, if a genomic region is active in human reference genome but inactive in a model genome, it is a LoA enhancer of the model genome. (B) Number of GoA and LoA enhancers in each cell type across the three model genomes as a fraction of total 1kb genomic regions that were scored. (C) Fraction of repurposed and *de novo* GoA enhancers across 67 cell types.

We next explored the genomic features of GoA and LoA enhancers. Examining the sequence conservation, we found that an average of 8% of these enhancers in the model chimp genome and an average of 9% in the 1000g or random genome across all cell types overlap genomic regions that are conserved across 30 primate species. This significantly exceeds the expected 5.6% overlap of the human reference genome with conserved elements (one-sample t-test p-value < 10^−20^). Additionally, we observed that the density of GWAS SNVs (Methods) is significantly lower compared with H3K27ac peaks in a cell-type (paired t-test p-value < 10^-40^, Figure S3), suggesting that frequent turnover reduces the biological impact of enhancers.

### Novel enhancer activity emerges predominantly from enhancer repurposing

We compared the emergence of novel enhancer activity from enhancer repurposing and *de novo* origins in our model genomes. GoA enhancers in each cell type were analyzed against a non-redundant dataset of H3K27ac peak regions from 111 biosamples (see Methods). GoA enhancers in a particular cell type that originate from enhancers currently active in other cell types were designated as repurposed, while those originating from non-enhancer sequences were dubbed *de novo* GoA enhancers. We found that 71-93% of the GoA enhancers were repurposed, while only a small fraction emerged as *de novo* enhancers (Figure 1C).

This suggests that repurposing existing enhancers is more common due to the sequence composition of transcription factor binding sites, which presents a low evolutionary barrier for the development of novel binding sites (Vierstra et al., 2014). Our results provide the first large-scale empirical evidence that enhancer repurposing is significantly more prevalent than *de novo* evolution of novel enhancer activity, and suggests a reduced likelihood of utilizing the entire sequence space when whole genome is used as a template. As expected, *de novo* GoA enhancers feature diminished sequence conservation compared to repurposed enhancers (Figure S4).

### Repurposed enhancers generate species and cell type specific gene expression patterns

We next investigated the role of repurposed enhancers in generating differential gene expression profiles. We focused on instances of enhancer repurposing in NPCs and hepatocytes, specifically analyzing 1,885 chimp NPC enhancers that are also predicted as human hepatocyte enhancers (NPC active enhancers of the model chimp genome are dubbed ‘chimp NPC enhancers’). These enhancers are predicted to be active in chimp NPCs but inactive in human NPCs, suggesting their role in regulating neurodevelopmental processes. As predicted, target genes of these enhancers exhibit elevated expression in chimp NPCs (Figure 2A). Notable genes with at least a three-fold higher expression in chimp NPCs include ADGRG6, GLIS3, and GALNT10, all essential for neurodevelopment. For example, ADGRG6 is a G-protein coupled receptor that regulates myelination and auditory system (Monk et al., 2011), and myelination is thought to play a role in the development of human cognition (Monk et al., 2009).

**Figure 2:**
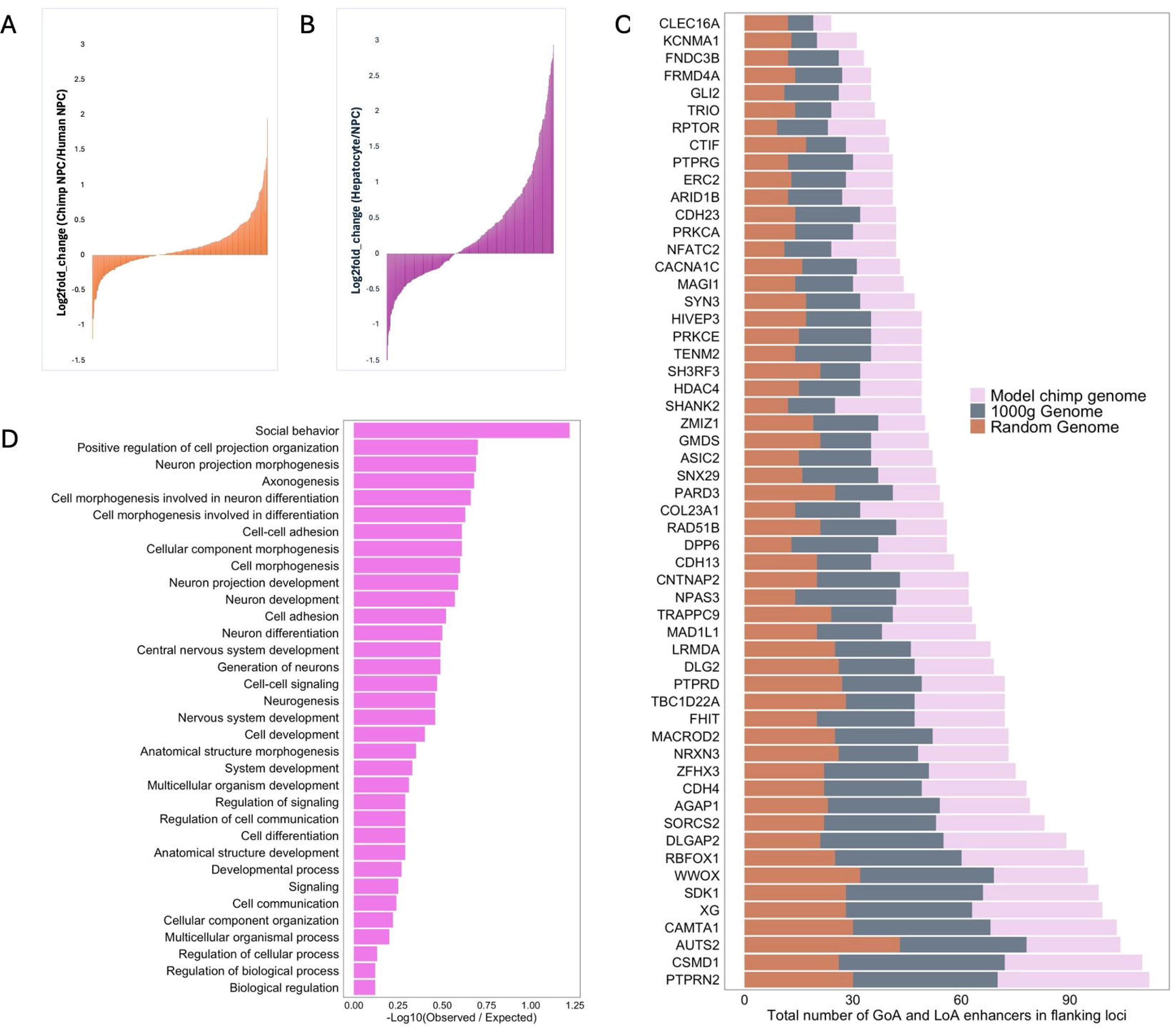
Target genes of GoA and LoA enhancers. (A) Log2fold change in chimp/human expression of genes in NPCs that within 100kb of enhancers active in chimp but not in human. (B) Log2fold change in Hepatocyte/NPC expression of genes that within 100kb of enhancers active in human hepatocyte but not in human NPCs. (C) Genes with highest turnover in their loci. The complete list of impacted genes is provided in Table S1. (D) Biological processes enriched in genes listed in (C). Terms with False Discovery Rate (FDR) < 10^-3^ are shown.

GALNT10 is a glycosyltransferase that’s upregulated in Schizophrenic patients (Voisey et al., 2017). We also identified NR2F2 as a differentially expressed gene with 15-fold higher expression in chimp NPCs. NR2F2 has been implicated in craniofacial feature divergence between the two species (Rada-Iglesias et al., 2013).

The loss of activity of these 1,885 enhancers in human NPCs is associated with a concurrent decrease in expression of flanking genes in NPCs when compared with human hepatocytes, where they still retain activity (Figure 2B). Among the target genes affected, ADGRG6, SLIT3, ATP8B1, and SLC13A5 are notable examples. These genes are not expressed in human NPCs but show 5-fold or higher expression levels in human hepatocytes. ADGRG6, although highly expressed in hepatocytes, lacks an established role in liver cell types. On the other hand, SLIT3, ATP8B1, and SLC13A5 have been demonstrated to function specifically in hepatocytes (Fu et al., 2024; Kumar et al., 2021; Naik et al., 2015). These findings demonstrate enhancer repurposing as a mechanism of cis- regulatory divergence that generates differential gene expression profiles between species and cell types.

### Genetic loci with high turnover are enriched for neurodevelopmental processes

Human-specific regulatory elements within primates have been shown to be biased towards embryonic and brain development (Whalen & Pollard, 2022; Won et al., 2019). While these human enhancer gains are often mediated by structural variants and transposable elements (Trizzino et al., 2017; Waterson et al., 2005), we were interested to explore the potential impact of SNVs in the human-specific traits. We thus turned our attention to genes that are targeted by GoA and LoA enhancers. We used genomic proximity to identify target genes as this approach has been shown to have a strong role in establishing enhancer promoter links (Gasperini et al., 2019). On average, 96% of the GoA and LoA enhancers from all cell types in the three model genomes are located within 100kb of at least one gene. A small set of genes, 269 in model chimp genome, 289 in 1000g genome and 366 in random genome were model genome exclusive targets of enhancer turnover (Table S2). The exclusive gene sets impacted in the 1000g genome are enriched for sensory perception and olfactory transduction (p_adj_ < 0.01, Table S3). This is consistent with several observations that have been documented in previous studies demonstrating the ongoing human specific loss of olfactory receptor genes (Gilad et al., 2003). Genes exclusively targeted in model chimp genome were only enriched for chromosome organization (Table S3). No enrichment was observed for exclusive genes of random genome.

Next, we obtained a set of 160 genes that were among the top 10 genetic loci with highest occurrence of GoA and LoA enhancers in each cell type, for the three model genomes (Figure 2C). This approach allowed us to pinpoint genes regulated by multiple enhancers that are prone to enhancer turnover. Interestingly, we obtained several genes common to the three model genomes. Notably, CNTNAP2, NPAS3 and AUTS2 have also previously been characterized as some of the genes with highest numbers of Human Accelerated Regions (HARs) in their regulatory loci (Kamm et al., 2013; Oksenberg & Ahituv, 2013). We found that AUTS2, along with PTPRN2, CAMTA1, XG and CSMD1 contain the highest number of GoA and LoA enhancers in their loci in all the three model genomes and are known for their roles in establishing memory, neuronal disorders and cancers (Chang et al., 2013; Ermis Akyuz & Bell, 2022; Huentelman et al., 2007; Lee, 2019). We investigated the annotation of these 160 genes through the Gene Ontology knowledgebase (The Gene Ontology Consortium et al., 2023) and found them to be enriched in social behavior, neuronal development and differentiation (Figure 2D). These findings are consistent with other studies which have shown human specific divergence of enhancers involved in early development, especially neurodevelopment (Mangan et al., 2022; Prabhakar et al., 2006). Since these genes are flanked by multiple enhancers, they are more prone to changes in enhancer activity across various mutational contexts represented by the three model genomes, making them more likely to play recurrent roles in neuropsychiatric diseases (Kamm et al., 2013).

We also hypothesized that simultaneous gain and loss of enhancer activity may create differences in the number of flanking enhancers leading to differential gene expression. We compared the differential expression of genes flanked by turnover loci in human and chimp NPCs with the net number of enhancers in the model chimp genome (calculated from the difference in counts of GoA and LoA enhancers) and found that gene expression is proportional to the number of flanking enhancers (Figure S5). Thus, enhancer turnover generates species-specific differences in the number of enhancers which further contributes to differential gene expression profiles. The maximum difference (four in the model chimp genome and one in the human reference genome) occurred in the locus of Syn3, that has elevated expression in chimp NPCs. Syn3 belongs to the Synapsin family of proteins known to regulate dopaminergic neuron development and has been implicated in Parkinson’s disease due to its interaction with alpha-synuclein and promoting aggregation (Zaltieri et al., 2015). Moreover, the intronic region of SYN3 which undergoes enhancer turnover in the model chimp genome, is enriched for variants associated with several neurological disorders (Longhena et al., 2021; Pizzollo et al., 2022).

### Enhancer turnover between human and chimp generates species specific transcription factor binding site profiles

Enhancer gain and loss of activity typically involves the introduction, loss, strengthening, or weakening of transcription factor binding sites (TFBSs). We focused on NPCs as a case study to investigate brain-specific changes, given that neuronal maturation in the brain is a significant factor in human-chimp divergence (Somel et al., 2009). We examined the TFs whose binding sites are enriched in LoA enhancers of NPCs in the model chimp genome, which are regions active in humans but not in model chimp genome. Compared with H3K27ac regions of human NPCs, these LoA enhancers are enriched for binding sites of general transcription regulators (Figure 3A). Importantly, they are depleted in neuronal lineage determining TFs such as SOX2, PHOX2A/B, OLIG1, etc. (Panman et al., 2011), indicating that core regulatory networks determining cell fate and identity remain intact. These findings align with previous studies demonstrating the conservation of core developmental networks during enhancer evolution among mammals (Andrews et al., 2023; Stergachis et al., 2014).

**Figure 3:**
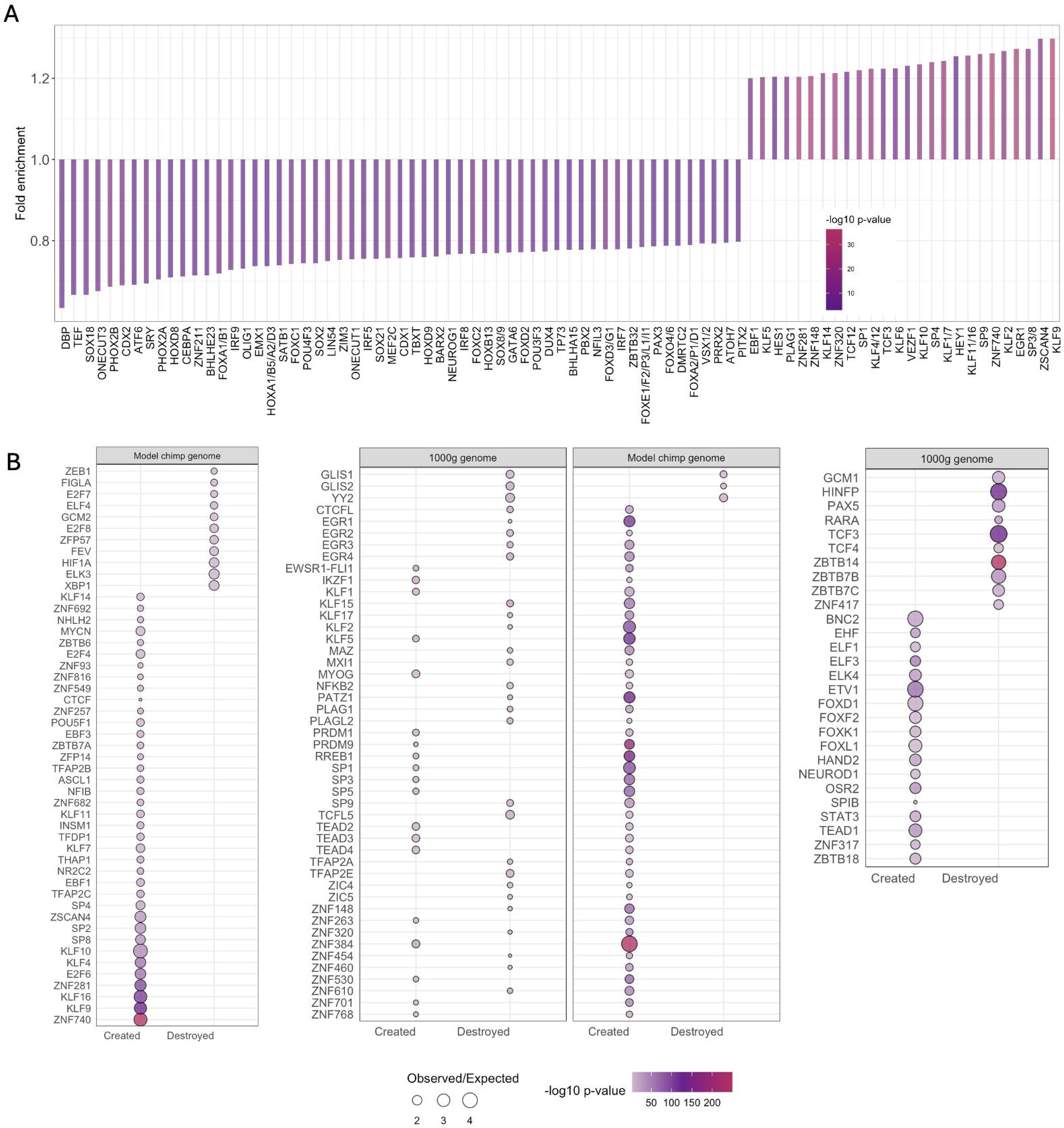
TFBS profiles of GoA and LoA enhancers. (A) Enrichment of TFBSs in LoA enhancers compared with H3K27ac regions in NPCs. P-values are calculated using Fisher’s exact test. (B) TFBSs that are created or destroyed at an accelerated rate. Binomial p-values obtained from observed/expected rate of creation or destruction for each TFBS (see Methods).

Next, we investigated TFBSs that may have changed at an accelerated rate among chimp GoA enhancers leading to their differential abundance in model chimp genome. We hypothesized that these changes, regardless of the TFBS enrichment in GoA enhancers of the model chimp genome, could be predictive of species-specific enhancer activity. Our hypothesis was driven by a previous observation where differential abundance of TF footprints within HARs was predictive of differential reporter assay activity between human and chimp NPCs (Whalen et al., 2023). To investigate this, we computed the observed rates of TFBS creation or destruction within turnover loci, using a background of motifs introduced by 100 permutations of random mutations. We compared these rates to the actual changes in the model chimp genome. TFBSs with accelerated changes were identified based on a binomial test assessing the significance of observed and expected rates of creation or destruction (Figure 3B). Among the TFs showing highest observed/expected ratio of binding site creation in NPCs of model chimp genome were ZNF384, ZNF740, PRDM9, KLF2/10 and SP1. PRDM9 acts as a recombination regulator and has been indicated to be a significant factor leading to genomic differences between the two species [24]. ZNF740 sites have also been shown to be more abundant in chimp NPC enhancers compared to human NPCs (Pizzollo et al., 2022). Conversely, we observed an accelerated rate of destruction of binding sites for TFs like HIF1A, BARHL1, and ZBED1. HIF1A regulates myelination and influences human cognition (Miller et al., 2012). Based on these findings, we propose that TFBSs undergoing accelerated motif creation and destruction in the model chimp genome serve as potential signatures of chimp and human NPC enhancers, respectively, and drive species-specific enhancer activity.

We expanded our analysis to the 1000g genome to identify differences in evolutionarily constrained and population-specific patterns of TFBS evolution. Several zinc finger TFs with accelerated creation rate in the model chimp genome also undergo accelerated change in the 1000g genome, indicating ongoing evolution within the human lineage (Figure 3B). Zinc finger TFs have been known as one of the fastest evolving classes of TFs among primates (Emerson & Thomas, 2009). Binding sites of TFs such as the TEAD factors, which are constituents of the Hippo pathway that has a central role in several types of cancer (Lamar et al., 2012), are created in both model genomes, implying a fluid nature of TEAD-dependent enhancer activity. We also observed another category of TFs whose binding sites show an accelerated rate of creation in the model chimp genome but are destroyed in the 1000g genome. These include PATZ1 that has a role in maintaining stem-cell properties in NPCs (M. Huang et al., 2024), neuroplasticity determining EGR4 is involved in learning and memory (L. Li et al., 2005) and ZIC proteins that regulate neuronal maturation (Minto et al., 2024) . Collectively, their accelerated evolution in both genomes is suggests a continuum in the evolution of the human brain.

### Cellular repurposing of enhancers suggests the role of non-cognate cell types in the biological impact of disease variants

Having established the role of enhancer repurposing in the evolution of species-specific gene expression profiles, we next explored their functional significance arising from variation within the human population. Trait associated variants are enriched in active enhancers defined by H3K27ac peaks (Hou et al., 2023). We used hepatocytes as a testbed and assessed the enrichment of GWAS traits in H3K27ac regions that undergo turnover in the 1000g genome.

First, we observed that the number of enhancers gaining or losing activity in hepatocytes is tenfold greater than those that remain unchanged in the 1000g genome. Consequently, very few but hepatocyte specific traits were associated with these regions (Figure 4A). Interestingly, both GoA and LoA enhancers contain several traits unrelated to hepatocytes. For instance, the GoA enhancers are enriched for traits associated with coronary artery disease (CAD), a condition primarily affecting the heart. However, CAD often presents with metabolic abnormalities linked to the liver, as established in previous studies (Selvarajan et al., 2021). CAD-enriched hepatocyte GoA enhancers have higher H3K27ac intensity in aorta, its cognate cell type (Figure 4B). Additionally, flanking genes of these loci have an elevated expression in aorta compared with hepatocytes (Figure 4C). These genes include JCAD, CFDP1, NT5C2, and FN1, all of which are known to be implicated in cardiovascular diseases. Our results indicate that variants in common enhancers of hepatocytes and aorta may contribute to the risk of CAD by increasing expression of their target genes in heart cells. These results are supported by recent MPRA studies confirming this effect in the genetic loci of CFDP1 and NT5C2 (Selvarajan et al., 2021).

**Figure 4:**
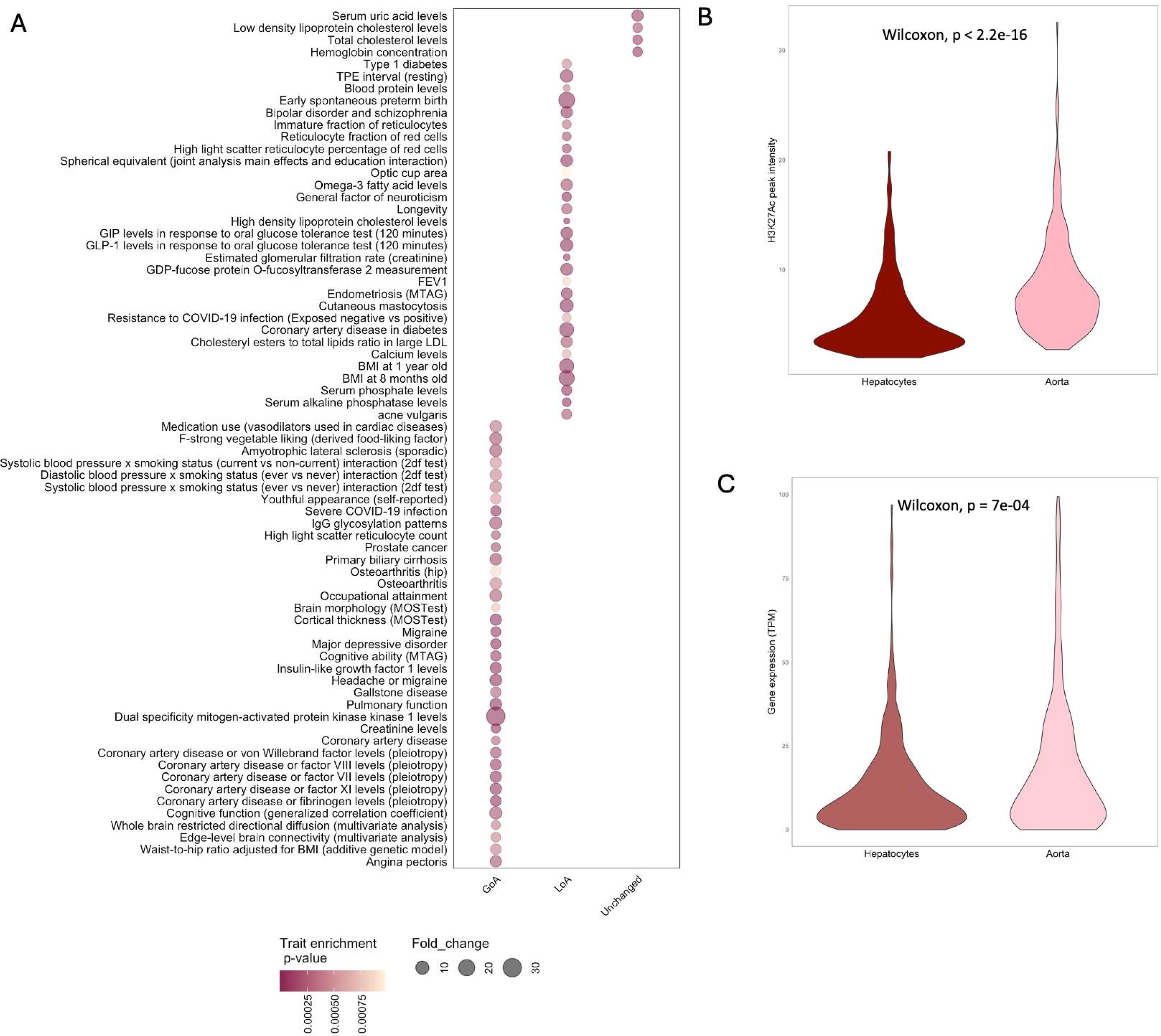
GoA and LoA enhancers of hepatocytes are enriched for non-cognate GWAS traits. (A) GWAS traits enriched in GoA, LoA and unchanged loci in H3k27ac peak regions of hepatocytes. Traits with at least 40 GWAS + LD SNVs were analyzed for enrichment. P-values calculated using Fisher’s exact test and traits that are enriched with p-value < 10^-3^ are shown. (B) Comparison of H3K27ac signal of enhancers enriched in CAD associated traits. (C) Comparison of expression of target genes within 100kb of enhancers analyzed in (B).

We extended these conclusions in assessing the locus enriched for variants of creatinine levels, another non-hepatocyte kidney associated trait that’s enriched in GoA enhancers of hepatocytes. One of the flanking genes of this locus is SLC22A2 that encodes for a creatinine transporter with restricted expression in kidney (Kumar et al., 2021). However, gain of activity in hepatocytes may lead to ectopic expression of the transporter, potentially resulting in increased uptake of creatinine in hepatocytes. This phenomenon could explain creatinine-induced hepatic damage observed as a phenotype of altered creatinine levels (Liu et al., 2022).

Although these findings are preliminary and need experimental confirmation, they offer strong evidence for investigating the role of causal variants that result in enhancer repurposing and involve non-cognate cell types in complex disease phenotypes.

### Random variants in the human genome identify vulnerable enhancers which are more likely to lead to enhancer turnover

Mutational robustness in enhancers is a mechanism that contributes to phenotypic stability in biological processes, particularly in regulating genes crucial for development. Enhancer activity can remain resilient to single nucleotide changes or even multiple mutations, resulting in both mutationally robust and vulnerable enhancers (S. Li et al., 2023). These categories together form a regulatory complement that allows for evolutionary adaptation while maintaining robustness in biological processes.

We aimed to explore the features of these two categories within enhancers of a cell type containing H3K27ac peaks. We thus expanded our approach to identify robust and vulnerable enhancers in six cellular contexts using predictions from ten model random genomes. Specifically, we scored ten model random genomes to identify robust enhancers with conserved activity across all model random genomes and vulnerable enhancers that lose activity in one or more of them. We observed variations in the proportions of robust and vulnerable enhancers across the six cell types. Notably, NPCs exhibit the lowest fraction of vulnerable enhancers, which aligns with expectations for a developmental cell line (Figure 5A). Additionally, robust enhancers showed significantly stronger evolutionarily constraint (Wilcoxon p-value < 10^-8^, Figure 5B) and higher H3K27ac peak intensities indicative of stronger enhancer activity compared to vulnerable enhancers, except for thymus and mesenchymal stem cells (Wilcoxon p-value < 10^-3^, Figure 5C). We also tested for differential enrichment of GWAS SNVs in the two categories of enhancers but found the differences to be insignificant overall.

**Figure 5:**
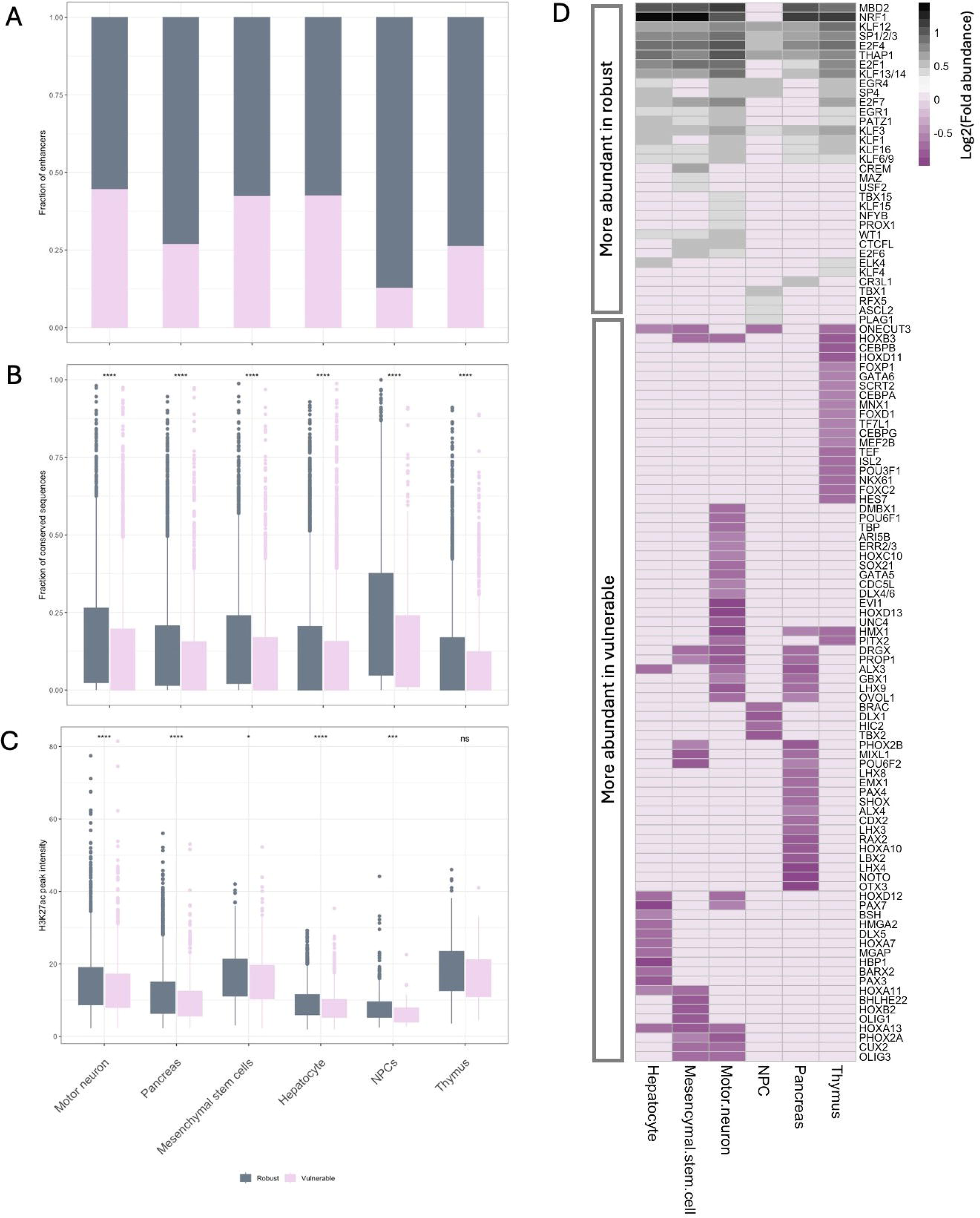
Features of robust and vulnerable enhancers in six cell types. (A) Fraction of robust and vulnerable enhancers. (B) Fraction of conserved elements in each category. (C) Comparison of H3K27ac peak intensities. (D) Differential abundance (robust/vulnerable) of TFBSs. A complete list of all TFs whose binding sites are differentially abundant in the two categories is provided in Table S4. ns: p > 0.05, *: p <= 0.05,*\*\*: p <= 0.01,***: p <= 0.001*

In parallel, we conducted an analysis on the differential abundance of TFBSs that distinguish the two categories of enhancers. Overall, robust enhancers across all cell types exhibited higher abundance of general transcription regulators and other globally expressed TFs such as the Specificity Proteins, Early Growth Response (EGR) factors, chromatin remodelers, and cell cycle regulators such as TFDP1 and ZBTB33 (Figure 5D, Table S4). In contrast, vulnerable enhancers were characterized by higher abundance of cell type-specific TFs such as SOX2, POU2F1, MEF2A of NPCs, CEBPA, HNF4A, etc., of hepatocytes, and CUX2 of motor neurons (Figure 5D, Table S4). These observations highlight vulnerable enhancers as hotspots for new enhancer activity, with the potential to develop cell type- or species-specific regulatory characteristics.

## 3. Discussion

Primate lineage has seen unprecedented changes in evolutionarily constrained regions of the genome (Ovcharenko, 2008). Recent research has demonstrated that HARs and human specific structural variants have a significantly higher association with TADs that are enriched in differentially expressed genes between the two species. As a result of target gene rewiring, HARs were thus subjected to different selection pressures (Keough et al., 2023). HARs are regions which have evolved rapidly in the human lineage, particularly near brain enhancers, diverging from regions that were perfectly conserved for millions of years (Whalen & Pollard, 2022). HARS and human gained enhancers (HGEs) are among the SNVs that contribute to approximately 1.2% of the observed differences between the human and chimp genome (Uebbing et al., 2021; Waterson et al., 2005). Thus, SNVs play a significant role in speciation events.

The availability of epigenomic profiles from various cell types enabled us to predict cell type-specific changes arising from genome-wide SNVs using DL models trained on open-chromatin and H3K27ac peaks of 67 cell types. Our results provide empirical evidence supporting enhancer repurposing as the primary source of regulatory innovation between closely related species. Moreover, through different mutational landscapes we find that enhancer innovation is limited to only ∼6% of the genomic landscape. A previous study comparing human and mouse DNAse Hypersensitivity Sites (DHS) regions reported that approximately 35% of the shared sites have undergone repurposing from one cell type to another between the two species (Vierstra et al., 2014). In contrast, we observe a significantly higher average of over 80% enhancer repurposing across all cell types between the human reference and the model chimp genome, reflecting the close evolutionary relationship between humans and chimpanzees. Importantly, these changes do not appear to disrupt the functional complement of enhancers within a cell type or the binding sites of lineage-specific TFs, regardless of the model genome used for comparison.

Our results demonstrate that regulatory innovations induced by SNVs frequently manifest in the cell type-specific enhancers that have a higher potential for adaptive changes. However, in a comparison of human and marmoset H3K27ac profiles, many recently evolved human specific enhancers were found to exhibit high interindividual variability and were less likely to have a biological impact as opposed to functionally conserved enhancers that effectively stabilize gene expression (Berthelot et al., 2018; Castelijns et al., 2020). Thus, enhancers undergoing turnover in the three model genomes belong to this class of newly evolved enhancers that may have a smaller and cell-type specific impact on gene expression. However, these changes create a distinct TFBS footprints that drive species-specific regulatory activity (Whalen et al., 2023). A comprehensive knowledge of cell-type specific TF networks is crucial to understand how the distinct footprints translate changes in binding sites to species and cell-type specific changes in gene expression.

A comparison of TF footprints in human and mouse DHS sites in a previous study exposed a striking conservation in the trans-regulatory network of TFs despite significant turnover in TFBSs (Stergachis et al., 2014). Conserved TF regulatory motifs were found to be driven by ubiquitous general transcription regulators such as SP1, EGR1, CTCF, etc., that occupy driver positions of regulatory motifs. These driver TFs regulate the expression of cell type specific TFs. The abundance of general transcription regulators within robust enhancers suggests their role as driver elements. Furthermore, robust enhancers also exhibit higher H3K27ac intensities, indicating potentially higher enhancer activity and therefore, a stable binding of transcription complexes. Driver TFs binding to robust enhancers thus play a crucial role in regulating and stabilizing the expression of cell type-specific TFs. In contrast, the TFBS plasticity in vulnerable enhancers leads to abundance of cell-type specific TFBSs that collaborate with driver TFs to generate cell-type specific transcription networks. This combined action of robust and vulnerable enhancers thus preserves a trans-regulatory network while facilitating evolutionary innovation in cis-regulatory sequences driven by adaptation.

Comparative genomic studies provide insights into the evolution of gene expression and enable us to identify regulatory mechanisms that govern complex traits and diseases. However, this study has technical caveats that precludes a holistic understanding of combinatorial changes that drive gene expression. Our analysis is centered on enhancers based on collective evidence from various studies which demonstrate the role of enhancer divergence in shaping species specific features (Pizzollo et al., 2022; Prescott et al., 2015; Uebbing et al., 2021; Vierstra et al., 2014; Villar et al., 2015). However, it has been shown that a majority of species-specific differences, especially among the hominins, are also driven by structural variants and transposable elements (Choi & Lee, 2020; Trizzino et al., 2017). These elements have the potential to cause genome scale alterations that can rewire gene regulatory networks by modifying the three- dimensional genomic contacts. Other mechanisms include variants that alter chromatin accessibility and methylation profiles (Gokhman et al., 2020; Weiss et al., 2021), and trans regulatory factors such as changes in cellular composition that contribute to species’ divergence (Barr et al., 2023). Recently developed large language models such as Enformer have demonstrated efficiency in predicting changes in gene expression and outperform other DL models in capturing long range regulatory interactions (Avsec et al., 2021). Future experiments that utilize the predictive power of such models to quantify the regulatory impact of various epigenetic alterations in cell-type specific contexts can complement our findings to elucidate bona fide regulatory network perturbations that alter gene expression.

## 4. Methods

### Generating Model genomes

We quantified the single nucleotide substitutions between human and chimp by comparing the multiple alignment file of GRCh38 and PanTro6, the human and chimp reference genomes respectively, from UCSC (https://hgdownload.soe.ucsc.edu/goldenPath/panTro6/vsHg38/). Insertions, deletions and other structural variations were not considered. A total of 36,621,296 substitutions were obtained with differing base conversion rates (Table S5). Model chimp genome was generated by replacing chimp specific alleles in the human reference genome. For 1000g genome, variants from 1000g catalog were obtained from the VCF file of all dbSNP variants for GRCh38 build (GCF_000001405.40.gz) from https://ftp.ncbi.nih.gov/snp/latest_release/VCF/. The number of human-chimp substitutions, i.e., 36,621,296 positions were randomly chosen from the VCF file to generate the 1000g genome using GATK (Auwera & O’Connor, 2020). In addition, ten random genomes were generated by introducing random variants at 36,621,296 random positions but with conserved human-chimp base conversion rates (Table S5).

### Cell type specific DL models

We used a two phase TREDNet model developed in our lab for cell type specific enhancer prediction (Hudaiberdiev et al., 2023). The first phase of the model was pre-trained on 4560 genomic and epigenomic profiles, which included DHS, ATAC-Seq, Histone ChIP-Seq and and TF ChIP-Seq peaks from ENCODE v4 (Luo et al., 2020). The second CNN is trained on the output vector of first CNN (containing predictions for 4,560 features) using cell-type specific putative enhancers defined by DHS and H3K27ac peaks as input. These training datasets were generated using steps described below. Chromosomes 8 and 9 were held out for testing, chromosome 6 was used for validation and other autosomal chromosomes were used to build the second phase model. The area under the ROC and PRC curve for each of these models is listed in Table S6.

Open chromatin (DHS or ATAC-Seq) and H3K27ac profiles for 67 cell types were downloaded from ENCODE (Luo et al., 2020) (Table S7). Positive datasets were defined as 2 kb regions centered on overlaps between H3K27ac ChIP-seq peaks and chromatin accessibility peaks of each cell type, excluding coding sequences, promoter proximal regions (<2kb from TSS) and ENCODE blacklisted regions (Amemiya et al., 2019). The 67 cell types analyzed in this study are a subset of all ENCODE biosamples with positive dataset size of at least 10,000 putative enhancers. A 10-fold control dataset was generated for each cell type using randomly sampled 2kb fragments of the genome, excluding the positive dataset of that cell type and blacklisted regions.

For each 2kb fragment, the model generates an enhancer probability score. Thresholds for predicting active enhancers were set at a False Positive Rate (FPR) of 1% with a positive to control dataset in the ratio 1:10.

### Gain and loss of enhancer activity

We created a bed file of adjacent 1kb windows tiled on the human reference genome excluding centromeric, telomeric, coding and TSS proximal regions. These coordinates were flanked with 500 base pairs and scored using the cell type-specific TREDNet models. LoA enhancers were identified as regions that score above the threshold in the human reference genome and below the threshold in the model genome, while GoA enhancers were regions that score below the threshold in the reference genome and above in the model genome. Regions that scored above the threshold in both reference and model genomes were predicted as conserved enhancers.

### Enrichment analysis of TFBSs

We used command line FIMO (Grant et al., 2011) to identify potential binding sites by scanning vertebrate TF motifs in JASPAR (Rauluseviciute et al., 2024) and HOCOMOCO (Vorontsov et al., 2024) databases along the sequences. P-value of 10^-5^ was used as threshold for FIMO based motif scanning. Motif enrichment in LoA enhancers relative to H3K27ac peaks in NPCs (Figure 3A) was performed using Fisher’s exact test.

To analyze the rate of change of TFBSs in GoA enhancers of NPCs in the model chimp genome, observed rates were obtained by comparing the number of TFBSs in the coordinates of GoA enhancers between human reference genome and model chimp genome. Next, hundred distinct sets of these GoA enhancers were generated by introducing N random mutations in each set where N is the number of substitutions in the GoA enhancers between human reference and the model chimp genome. Expected rate was derived as the average of counts obtained for each motif from the 100 random permutations of GoA enhancers. Significance in rate of motif creation or destruction was performed using binomial test between observed and expected rates. The same process was repeated for GoA enhancers of 1000g genome.

Differential abundance of TFBSs in robust and vulnerable enhancers was calculated using the ratio of normalized count of binding sites of a TF in robust vs. vulnerable enhancers. The number of binding sites of all TFs in each category were normalized using the total number of non-overlapping binding sites identified by FIMO in that category of enhancers.

### Data and tools

The H3K27ac peaks used to validate TREDNet’s predictions of human and chimp NPC enhancer activity were downloaded from another study (Whalen et al., 2023) (GEO accession: GSE110760, N2 stage). The hg19 coordinates of these peak files (for both species) were converted to hg38 coordinates using liftOver (Raney et al., 2024). Orthologous H3K27ac peak regions between human and chimp were identified from peak centers overlapping by a minimum of 1 base pair. The same criterion was used to compare H3K27ac peak intensities shared between hepatocyte and aorta. Gene expression data for comparison between human cell types was downloaded from ENCODE. Gene expression profiles of human and chimp NPCs were obtained from another experiment (GEO accession: GSE127253). Raw counts were downloaded for replicates at day 14 for both species and normalized using DESeq2 (Love et al., 2014).

Evolutionary conservation of genomic regions was measured by their extent of overlap with phastCons elements conserved across 30 primates (https://hgdownload.soe.ucsc.edu/goldenPath/hg38/database/ phastConsElements30way.txt.gz). Density of GWAS SNPs in the loci of interest was computed using a dataset of 1.4 million SNVs downloaded from the NHGRI-GWAS catalog including SNVs in tight linkage disequilibrium (r^2^ >=0.8).

## Declarations Acknowledgements

This research was supported by the Intramural Research Program of the National Library of Medicine (NLM), National Institutes of Health. This work utilized the computational resources of the NIH HPC Biowulf cluster.

## Authors’ contributions

J.S. performed the computational analysis and analyzed the data. I.O. supervised computational work. J.S. prepared figures and tables. J.S. and I.O. wrote the manuscript.

## Availability of data and material

Please see the section “Data and tools” and supplementary tables.

## Ethics approval and consent to par5cipate

Not applicable.

## Consent for publication

Not applicable.

