## Supplementary material for "Regulatory plasticity of the human genome": Figure S*, Supplementary note *

**This PDF file includes:**

Supplementary notes 1 and 2

Supplementary Figures S1 to S7

**Supplementary Note 1**

Common variants between the model genomes are listed in Figure S2. To examine potential biases from these shared variants, we removed the ~2 million common variants between the model chimp and 1000g genome to create new model genomes. Due to a small overlap of variants, the new genomes did not show significant differences in turnover (Figure S6A, p > 0.001). Notably affected loci flank genes including include *GLI2* and *GLI3* (spinal cord development), *CELF4* (synaptic function regulation), and *CAMTA1* (memory) (Figure S6B). The presence of common variants between model chimp and 1000g genome highlights the dynamic nature of human-chimp substitutions, with some still present as polymorphisms in the human population.

**Supplementary Note 2**

Rare variants (MAF <= 0.01) constitute a major fraction of the 1000Genomes catalog. The ~36 million variants that were subsampled from the catalog to build the 1000g genome contain 31 million rare variants. In total, the 1000G catalog contains 13,731,885 common variants with MAF > 0.01. These common variants are significantly less conserved than the 36,621,296 variants that we had introduced in the 1000g genome (Mann-Whitney u-test p-value 10^-148^). To investigate possible discrepancies arising from rare and common variants, we incorporated the 13,731,885 common variants in the reference genome to generate a new 1000g genome (dubbed common variant 1000g genome) that does not contain any rare alleles and analyzed the resulting differences with the original 1000g genome. However, we found that exclusion of rare alleles had no significant impact on enhancer turnover across the 67 cell-types (Figure S7A). Moreover, the fraction of novel enhancers that emerge through repurposing or *de-novo* in the common variant 1000g genome are consistent with the fractions in 1000g genome (Figure S7B).

Further, we designed a new model chimp genome and a random genome containing the same number of randomly subsampled human-chimp substitutions and random variants, respectively, as the common variant 1000g genome and observed equivalent GoA and LoA enhancers across the three new model genomes (Figure S7C), consistent with our observations in the original model genomes (Figure 1B). Together, these results indicate that enhancer turnover is independent of allele frequency.

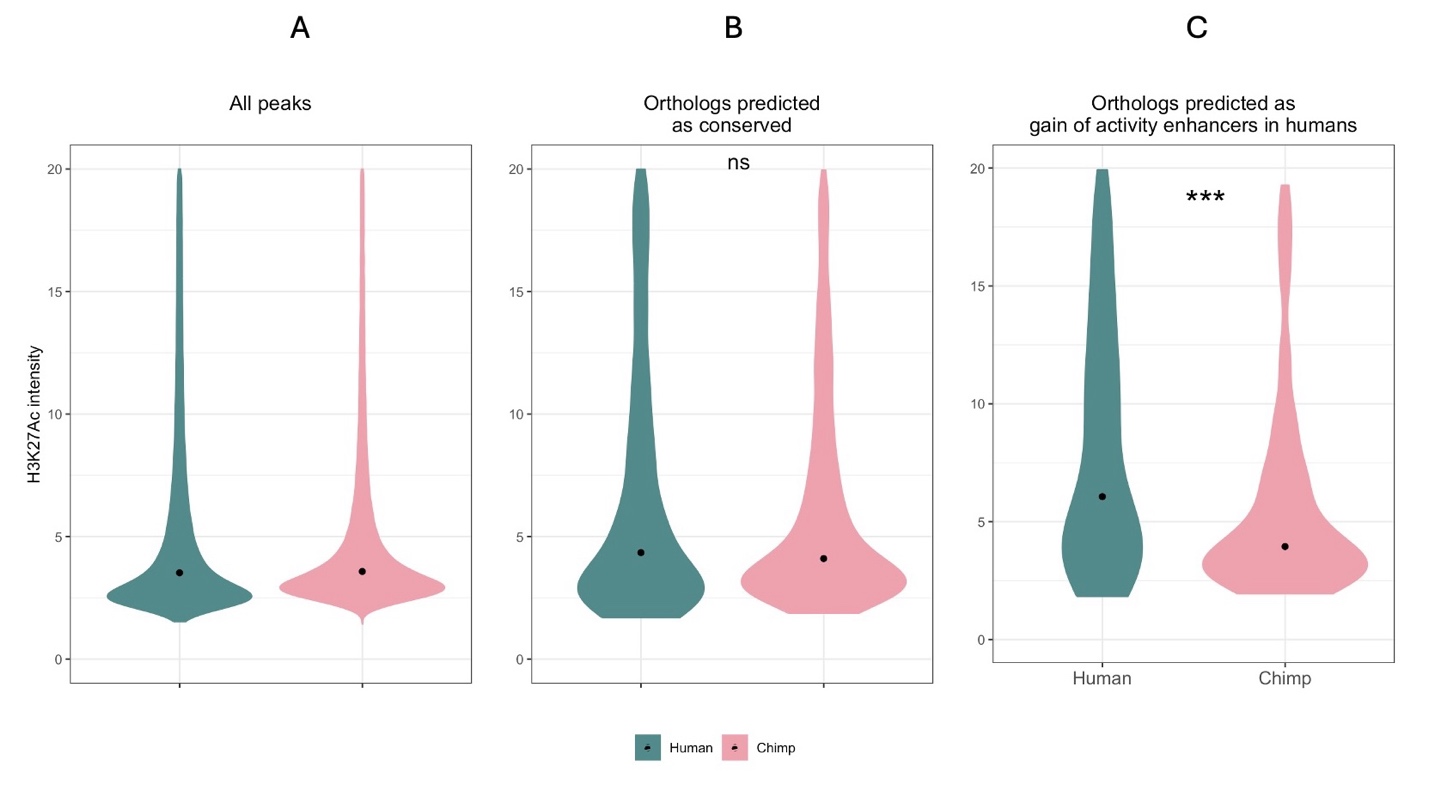

**Figure S1:** Comparison of H3K27ac peaks intensities of A) all human and chimp NPC enhancers and B) orthologous enhancers that are predicted as conserved and C) orthologous enhancers that are predicted to gain activity in humans. Orthologous regions were defined using criteria described in Methods*. ns: p > 0.05, ***: p <= 0.001*

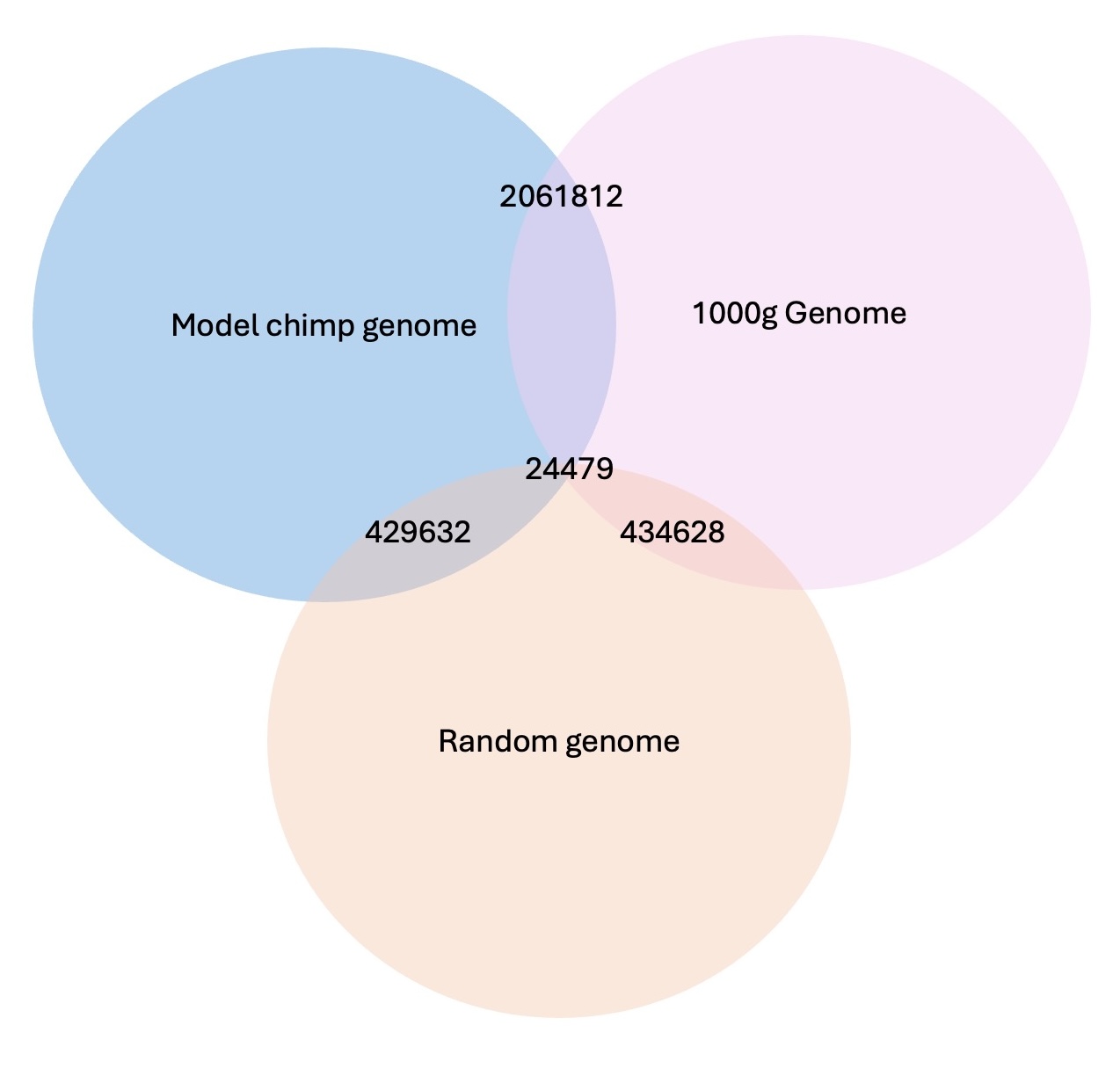

**Figure S2:** Number of common SNVs between the three model genomes. Total number of SNVs introduced in each model genome is 36,621,296. Overlaps are not scaled to actual numbers.

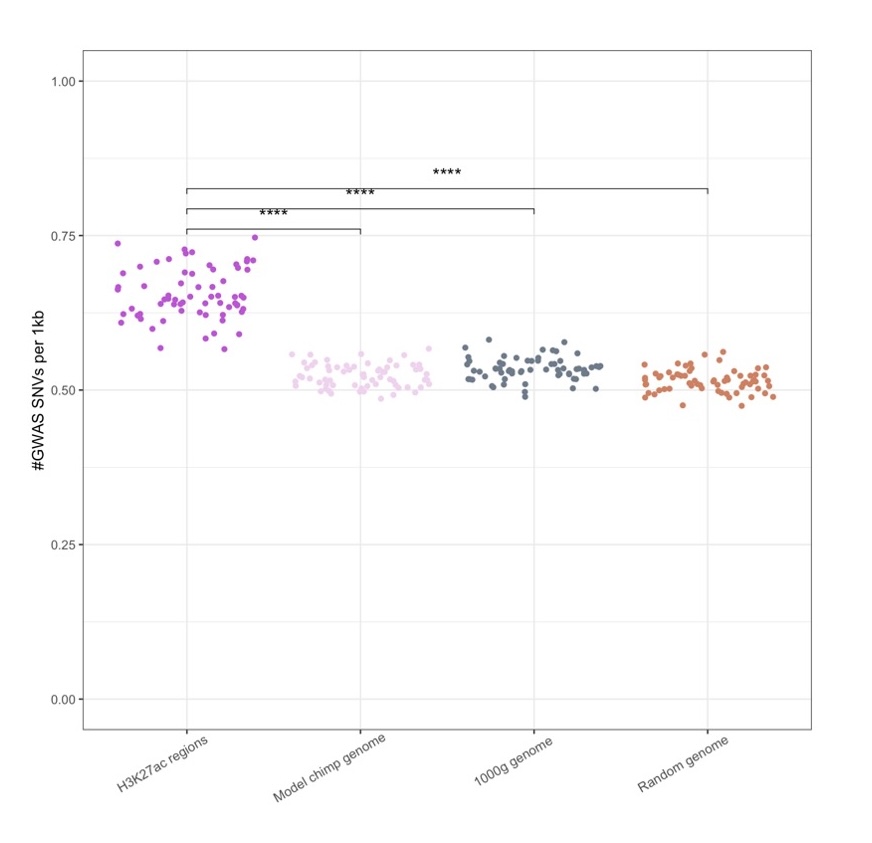

**Figure S3:** Density of GWAS SNVs (number of SNVs per 1kb) in the turnover loci of the three model genomes and in H3K27ac peak regions across cell types. ****: p <= 0.0001

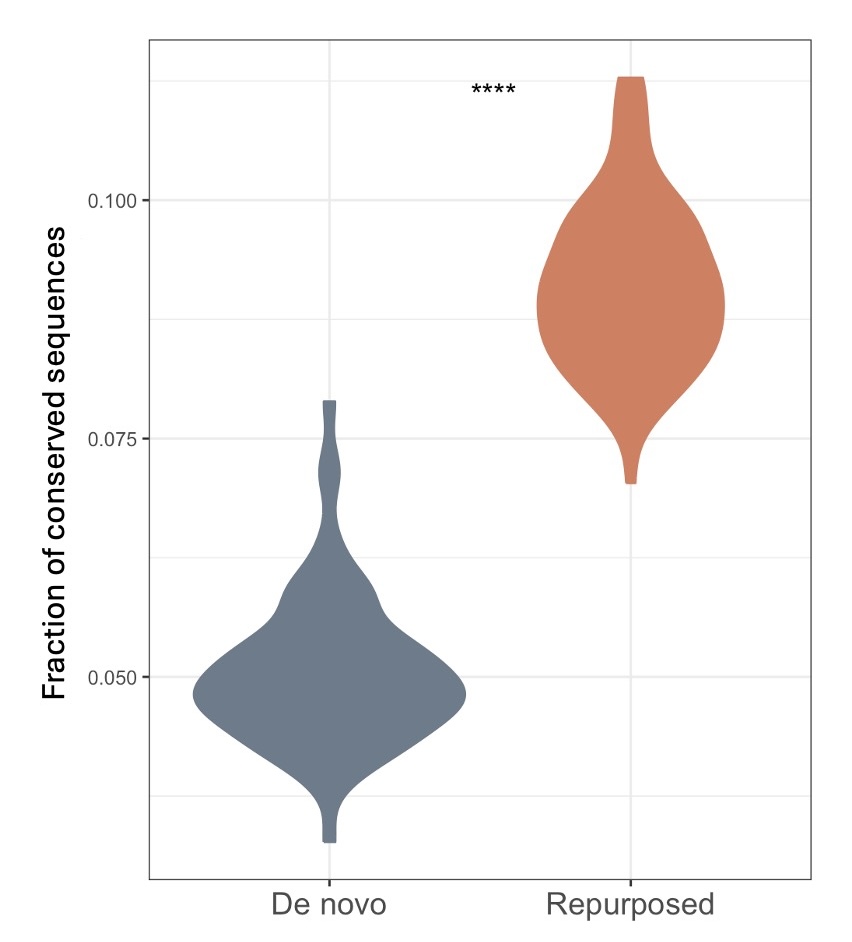

**Figure S4:** Fraction of conserved elements in GoA enhancers that emerge de novo or are repurposed, across all cell types. ****: p <= 0.0001

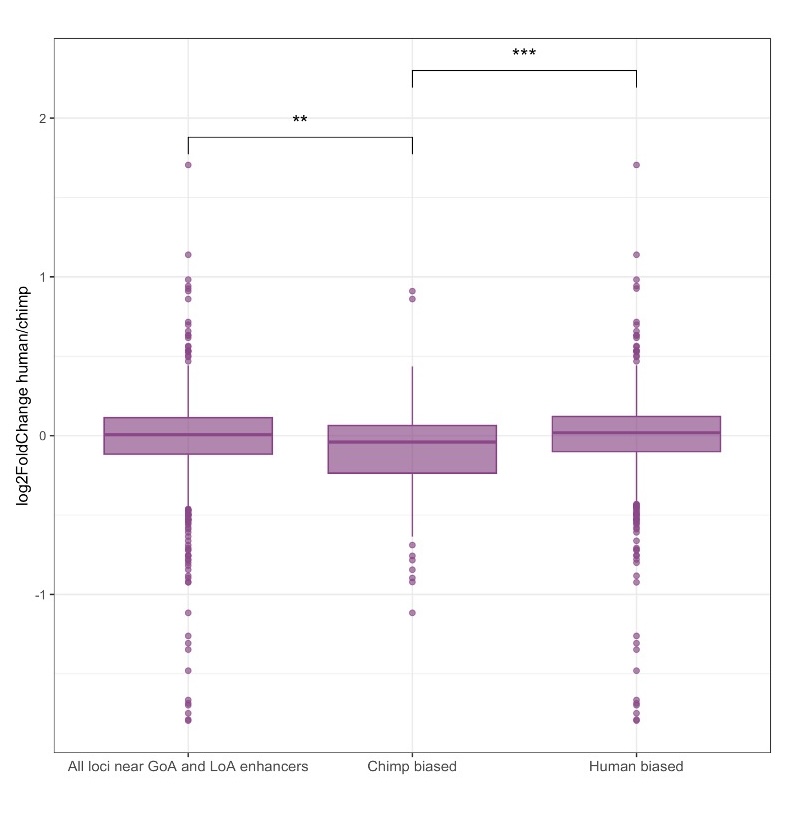

**Figure S5:** Log2fold change in human/chimp expression of genes in NPCs that are within 100kb of 1) all GoA and LoA loci of NPCs in model chimp genome, 2) higher number of GoA than LoA enhancers (chimp biased), and 3) higher number of LoA than GoA enhancers (human biased). **: p <= 0.01,***: p <= 0.001

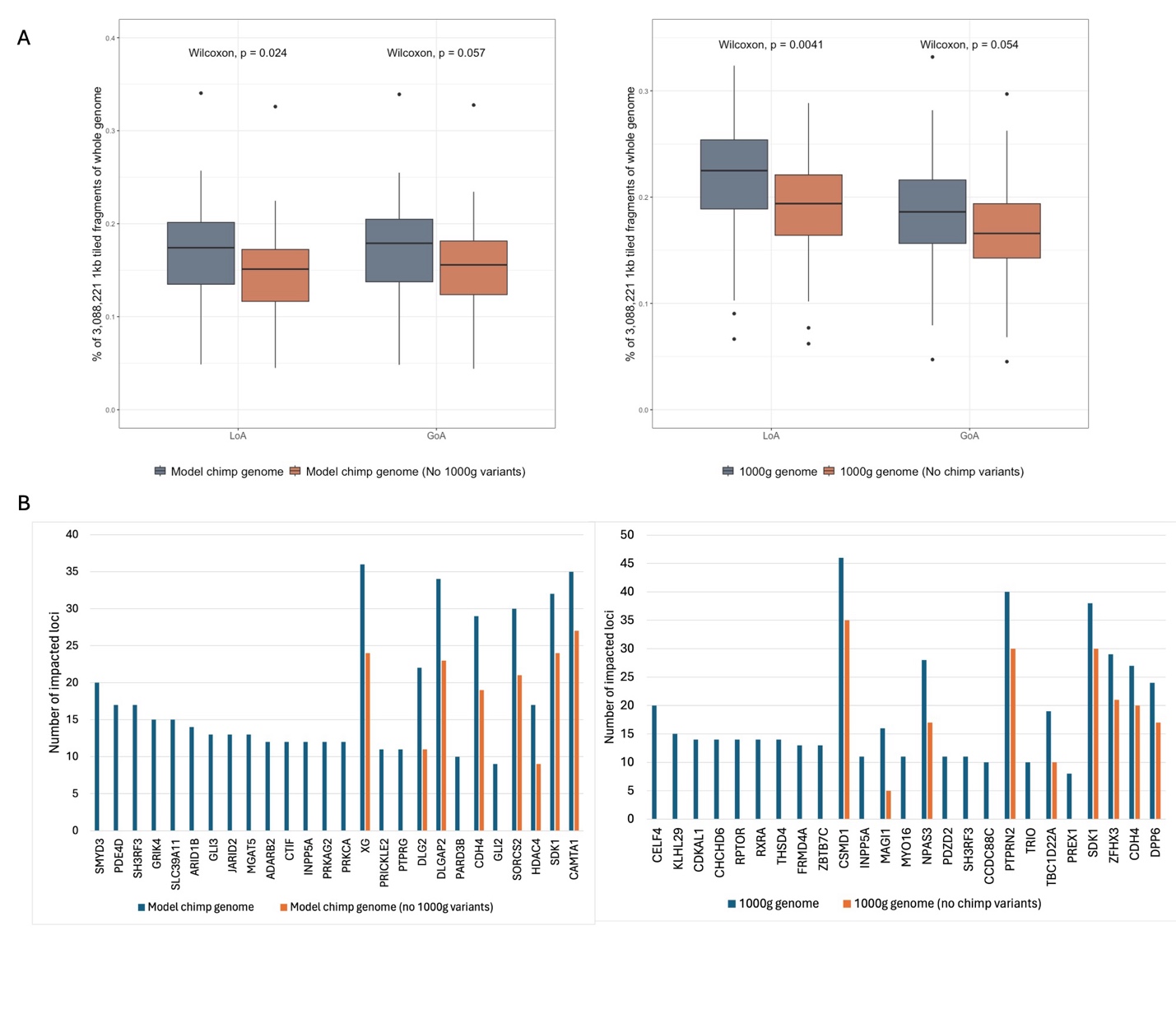

**Figure S6:** Impact of variants shared between the model chimp and 1000g genome. (A) A comparison of the fraction of GoA and LoA enhancers in the new model genomes without common variants with the fraction in original model genomes. (B) Top 25 genes with largest differences in their flanking enhancers across cell-types in the model chimp genome (left) and the 1000g genome (right).

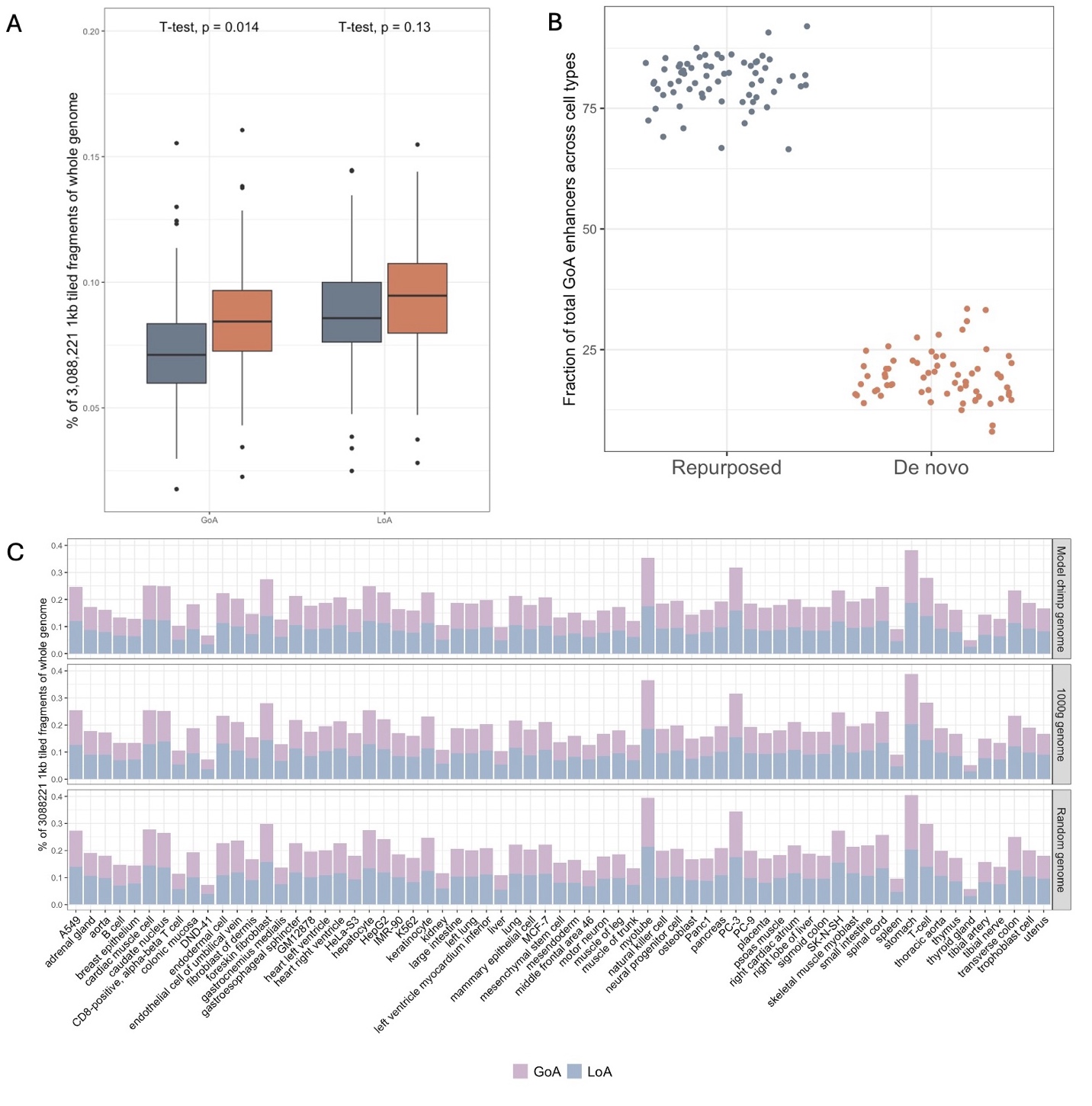

**Figure S7:** A comparison of 1000g genome and common variants 1000g genome. (A) Comparison of the number of GoA and LoA enhancers in the two model genomes. Turnover in 1000g was normalized with the ratio of number of common variants (13,731,885) and total number of variants used in original model genomes (36,621,296) (B) Fraction of repurposed and *de novo* GoA enhancers across 67 cell-types in the common variants 1000g genome. (C) Number of GoA and LoA enhancers in each cell type across the three model genomes as a fraction of all 1kb genomic regions.
